# *Paucimyces polynucleatus* gen. nov, sp. nov., a novel polycentric genus of anaerobic gut fungi from the feces of a wild blackbuck antelope

**DOI:** 10.1101/2021.03.04.433954

**Authors:** Radwa A. Hanafy, Noha H. Youssef, Mostafa Elshahed

## Abstract

The anaerobic gut fungi (AGF, phylum Neocallimastigomycota) reside in the alimentary tracts of herbivores. Multiple novel, yet-uncultured AGF taxa have recently been identified in culture-independent diversity surveys. Here, we report on the isolation and characterization of the first representative of the RH5 lineage from fecal samples of a wild blackbuck (Indian Antelope) from Sutton County, Texas, USA. The isolates displayed medium sized (2-4 mm) compact circular colonies on agar roll tubes and thin loose biofilm-like growth in liquid medium. Microscopic examination revealed monoflagellated zoospores and polycentric thalli with highly branched nucleated filamentous rhizomycelium, a growth pattern encountered in a minority of described AGF genera so far. The obtained isolates are characterized by formation of spherical vesicles at the hyphal tips from which multiple sporangia formed either directly on the spherical vesicles or at the end of sporangiophores. Phylogenetic analysis using the D1/D2 regions of the large ribosomal subunit (D/D2 LSU) and the ribosomal internal transcribed spacer 1 (ITS1) revealed sequence similarities of 93.5%, and 81.3%, respectively, to the closest cultured relatives (*Orpinomyces joyonii* strain D3A (D1/D2 LSU), and *Joblinomyces apicalis* strain GFH681 (ITS1). Substrate utilization experiments using the type strain (BB-3) demonstrated growth capabilities on a wide range of mono-, oligo-, and polysaccharides, including glucose, xylose, mannose, fructose, cellobiose, sucrose, maltose, trehalose, lactose, cellulose, xylan, starch, and raffinose. We propose accommodating these novel isolates in a new genus and species, for which the name *Paucimyces polynucleatus* is proposed. The type species is strain BB-3.

## Introduction

In the herbivorous gut, a diverse community of bacterial, archaeal, protozoan, and fungal species synergistically mediate the breakdown of plant biomass (Gruninger et al., 2014). Fungi in the herbivorous gut belong to a distinct fungal phylum (Neocallimastigomycota) and play a pivotal role in this process through mechanical and enzymatic means (Hess et al., 2020). Nineteen anaerobic gut fungal (AGF) genera have been characterized so far (Barr et al., 1989; Breton et al., 1990; Callaghan et al., 2015; Dagar et al., 2015; Gold et al., 1988; Hanafy et al., 2017; Hanafy et al., 2018; Hanafy et al., 2020b; Heath et al., 1983; Joshi et al., 2018; Ozkose et al., 2001; Stabel et al., 2020). However, culture-independent diversity surveys have identified representatives of multiple yet-uncultured AGF genera (Hanafy et al., 2020a; Kittelmann et al., 2012; Liggenstoffer et al., 2010). The amenability of such lineages to isolation is uncertain. The lack of cultured representatives could be a reflection of the complexity and difficulty in isolation and maintenance of these AGF lineages. Alternatively, it is possible that some yet-uncultured AGF taxa have complex nutritional requirements that are not satisfied in current media and isolation procedures.

Based on prior evidence (Hanafy et al., 2020a), we hypothesize that success in isolating a fungal taxon is directly proportional to its relative abundance within a specific sample. As such, targeting samples assessed to harbor a relatively large fraction of yet-uncultured taxa using culture-independent approaches should be prioritized in culture-based diversity efforts. During a recent culture-independent diversity survey of the AGF community in wild, zoo-housed, and domesticated herbivores in the US states of Oklahoma and Texas, we encountered several samples that harbored a high proportion of yet-uncultured genus-level clades of AGF (Hanafy et al., 2020a). We here report on the targeted isolation and detailed characterization of multiple strains belonging to one of these clades (lineage RH5). Morphological, microscopic, and phylogenetic characterization justifies proposing a novel genus and species to accommodate these isolates, for which the name *Paucimyces polynucleatus* is proposed.

## Materials and Methods

### Samples

Fresh fecal samples were collected from a wild blackbuck antelope during a hunting expedition in Sutton County, Texas, USA in April 2018. All hunters had the appropriate licenses, and animals were shot either on a private land with the owner’s approval or on public land during the hunting season. Samples were stored on ice and promptly (within 24 hours) transferred to the laboratory. Upon arrival, a portion of the sample was stored at -20° C.

### Isolation

A recent culture-independent survey of AGF diversity identified a wide range of novel yet-uncultured AGF lineages in fecal and rumen samples recovered from multiple herbivores in the states of Oklahoma and Texas (USA) (Hanafy et al., 2020a). Among the samples surveyed, a domesticated sheep and a wild blackbuck antelope showed a high relative abundance of the yet-uncultured lineage RH5 (96.2% and 52.4%, respectively) (Hanafy et al., 2020a). Due to the unavailability of sufficient feces from the domesticated sheep, isolation efforts were conducted only on the wild blackbuck antelope as previously described in (Hanafy et al., 2018; Stabel et al., 2020). Briefly AGF was enriched for 24 h at 39°C in rumen fluid (RF) media (Hanafy et al., 2017) amended with 0.5% cellobiose as a substrate. Enriched tubes were serially diluted in RF media supplemented with a (1:1) mixture of cellobiose and switchgrass. Antibiotics mixture (50 μg/mL penicillin, 20 μg/mL streptomycin, and 50 μg/mL chloramphenicol) was added to inhibit bacterial growth. Dilutions showing visible signs of fungal growth such as clumping and floating of the switchgrass, and/or production of gas bubbles were used for colony isolation using the roll tube procedure (Hungate, 1969). Purity of the obtained cultures was ensured by conducting three rounds of roll tubing and colony picking. Isolates were maintained by bi-weekly sub-culturing into cellobiose containing RF media. Long-term storage of the obtained isolates was conducted by surface inoculation on RF-cellobiose agar medium as previously described in (Calkins et al., 2016).

### Morphological and microscopic characterization

Three-day old colonies and liquid cultures were examined to describe the isolate’s growth pattern on solid and liquid media, respectively. Both light and scanning electron microscopy were utilized to examine different fungal structures at various stages of growth. For light microscopy, fungal biomass was stained with lactophenol cotton blue and examined using an Olympus BX51 microscope (Olympus, Center Valley, Pennsylvania) equipped with a DP71 digital camera (Olympus). Nuclei localization was examined by staining the samples with 4, 6′ diamidino-2-phenylindole (DAPI at final concentration of 10 μg/ml), followed by incubation in the dark for 10 min at room temperature. Treated samples were examined with LSM 980 confocal microscope with Airyscan 2 (Carl Zeiss AG, Oberkochen, Germany). Sample preparation and fixation for scanning electron microscopy was conducted as previously described (Hanafy et al., 2017; Hanafy et al., 2018). Samples were examined on a FEI Quanta 600 scanning electron microscope (FEI Technologies Inc., Oregon, United States).

### Substrate utilization

The substrate utilization capabilities of the type strain BB3 were assessed by using a rumen-fluid basal medium with no carbon source as previously described (Hanafy et al., 2017). Growth and viability of a 10% inoculum was compared to a substrate-free medium. Twenty-four different substrates were tested at a final concentration of 0.5% w/v (Table 1). The ability of strain BB3 to utilize a specific substrate was considered positive when the tested substrate supports the culture viability after three successive sub-culturing events.

**Table 1.** Substrate utilization pattern of *Paucimyces polynucleatus* strain BB-3.

| Substrate |  | Growth<sup>a</sup> |
| --- | --- | --- |
| Polysaccharides | Cellulose | + |
|  | Xylan | + |
|  | Starch | + |
|  | Raffinose | + |
|  | Inulin | - |
|  | Poly-galacturonate | - |
|  | Chitin | - |
|  | Alginate | - |
|  | Pectin | - |
| Disaccharides | Cellobiose | + |
|  | Succrose | + |
|  | Maltose | + |
|  | Trehalose | + |
|  | Lactose | + |
| Monosaccharides | Glucose | + |
|  | Xylose | + |
|  | Mannose | + |
|  | Fructose | + |
|  | Glucuronic acid | - |
|  | Arabinose | - |
|  | Ribose | - |
|  | Galactose | - |
| Peptides | Peptone | - |
|  | Tryptone | - |

a: +, Growth was observed following three consecutive subcultures; -, No growth was observed with the carbon source.

### Phylogenetic analysis and ecological distribution

Fungal biomass was harvested from actively growing 3-day old cultures and ground in liquid nitrogen. DNA was extracted from the ground biomass using DNeasy PowerPlant Pro Kit (Qiagen Corp., Germantown, MD, USA) according to the manufacturer’s instructions. The ITS1, 5.8S rRNA, ITS2 and D1/D2 region of the LSU rRNA was amplified using the primers ITS5 (5’ -GGAAGTAAAAGTCGTAACAAGG-3’) and NL4 (5’-TCAACATCCTAAGCGTAGGTA-3’) as described previously (Wang et al., 2017). Amplicons were purified using PureLink® PCR Purification Kit (ThermoFisher Scientific, Waltham, Massachusetts), and cloned using a TOPO-TA cloning vector according to the manufacturer’s instructions (Life Technologies®, Carlsbad, CA). Three clones were Sanger-sequenced at the Oklahoma State University DNA sequencing core facility. Regions corresponding to the ITS1 and D1/D2 LSU regions from the obtained amplicons were aligned to reference ITS1 and D1/D2 LSU sequences using MAFFT aligner (Katoh et al., 2019). Maximum likelihood phylogenetic trees were constructed in FastTree using *Chytriomyces* sp. WB235A isolate AFTOL-ID 1536 as an outgroup. Bootstrap values were calculated on the basis of 100 replicates.

To evaluate the ecological distribution of this novel lineage, we queried the ITS-1 sequences of isolates obtained from this study against GenBank nr (non-redundant) database using BLASTn and modified the output to display 5000 instead of the default 100 aligned sequences. Sequence similarity cutoff of 95% was used to filter the BlASTn output. The phylogenetic position of sequences with significant similarity (≥95%) was evaluated by inserting into ITS-1 reference phylogenetic trees.

### Data and culture accession

Sequences generated in this study are deposited in GenBank under accession numbers MW694896-MW694898. Cultures are available at Oklahoma State University, Department of Microbiology and Molecular Genetics culture collection. Genus and species information has been deposited in Mycobank under the ID number MB838953 and MB838954, respectively.

## Results

### Isolation

Multiple colony morphologies were obtained in roll tubes derived from enrichments of the feces of a wild blackbuck antelope. One colony type showed little morphological resemblance to currently described taxa, and its distinctness was confirmed by microscopic and phylogenetic analysis (see below). Four isolates (BB-12, BB-14, BB-2, BB-3) were examined, and all showed identical morphological and microscopic attributes. One isolate (strain BB3) was chosen as the type strain for detailed characterization.

### Morphology

On solid media, strain BB-3 formed white compact circular uniform colonies that lacked a darker central core of sporangial structures, often observed with monocentric AGF genera (Figure 1a). Colony size ranged from 2-4 mm. In liquid media, strain BB-3 forms a loose thin white biofilm-like growth (Figure 1b).

**Figure 1.**
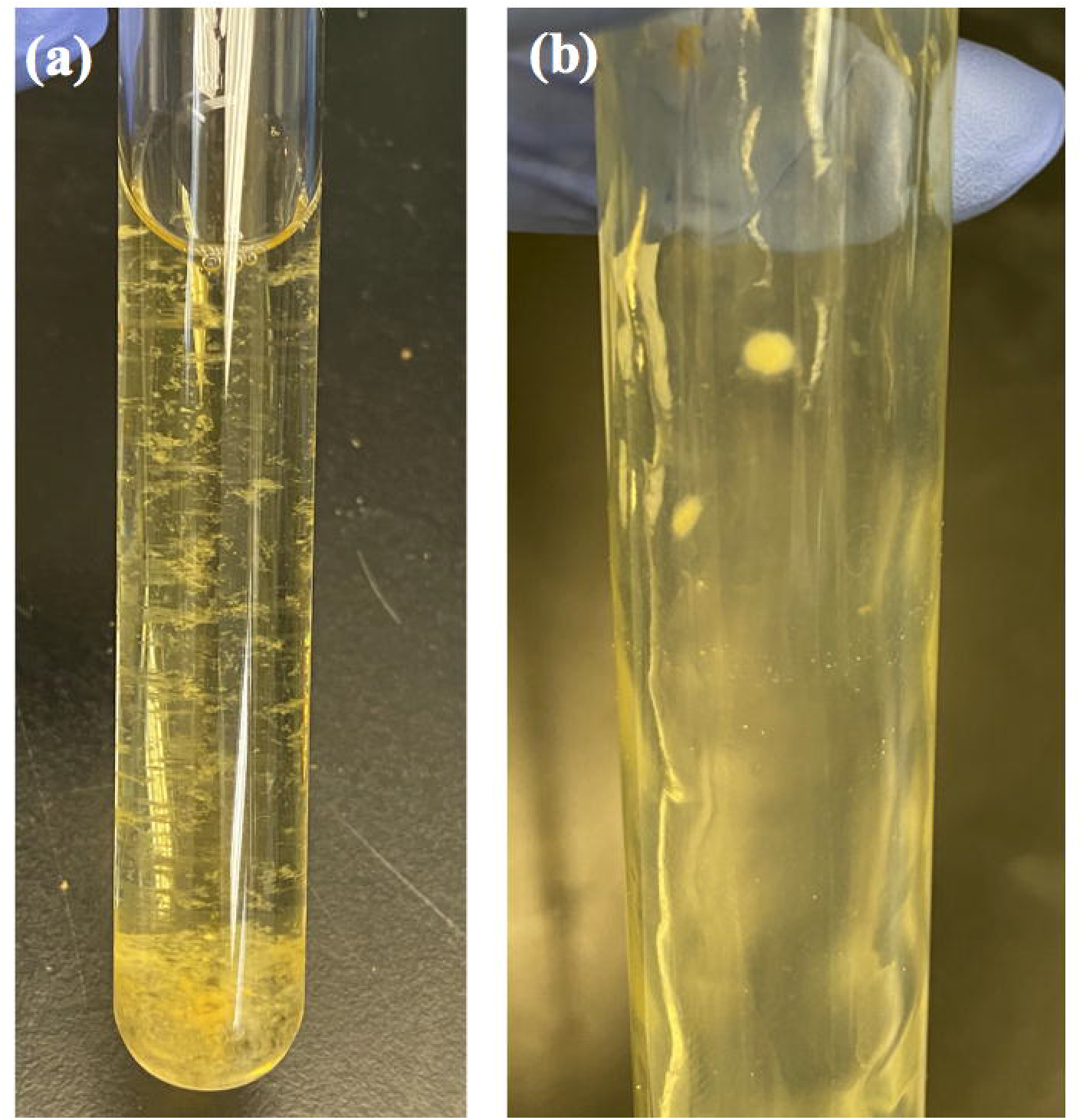
Macroscopic features of *Paucimyces polynucleatus* strain BB-3. (a) Thin and loose fungal biofilm-like growth in liquid cellobiose rumen fluid medium. (b) White, circular compact colony on cellobiose agar roll tube.

### Microscopic features

Strain BB-3 produces globose zoospores (Figure 2a-b), with an average diameter of 7.5 μm (range: 6-10 μm). The majority of zoospores were monoflagllated (Figure 2a), although biflagellated zoospores were occasionally observed (Figure 2b). Flagellum length ranged between 15-30 μm. Upon germination, zoospores contents migrated into the germ tube and the remaining empty zoospore cyst had no further function in the thallus development. This is in contrast to the zoospore cyst of monocentric genera that either enlarges into sporangia or develops a sporangiophore with a sporangium at the end (Ho and Barr, 1995). The germ tube eventually germinated to produce extensively branched polycentric thalli with nucleated filamentous rhizomycelium. The nucleated rhizomycelium produced multiple sporangia, giving rise to a polycentric thallus of indeterminate length (Figure 2c-f). During early thallus development, the hyphal tips started to swell forming spherical vesicles (Figure 2g), from which multiple sporangiophores arose (Figure 2h-i). Each sporangiophore had a single sporangium at its end (Figure 2c, d, i-k). In many cases, sporangia developed directly on the spherical vesicles without sporangiophores (Figure 2l-m). In rare occasions, strain BB-3 produced thalli with single sporangia (Figure 2n-o). Sporangia were mainly ovoid and ranged in size between 15–90 µm L x 10–55 µm W (Figure 2i-o). Upon maturity, basal walls were formed to separate the mature sporangia from the sporangiophores (Figure 2j-k). Old cultures appeared to progressively lose the ability to produce sporangia and only produced sporangiophores initials (Figure 2l), a distinct feature that was observed in old *Orpinomyces* cultures (Ho and Barr, 1995). Zoospores were released through a wide apical pore at the top of the sporangia, with the sporangial wall staying intact after the discharge (Figure 2r). Similar to the majority of polycentric AGF genera, strain BB-3 culture lost its zoosporogenesis ability due to frequent sub-culturing and started to produce sterile sporangia that did not differentiate into zoospores.

**Figure 2.**
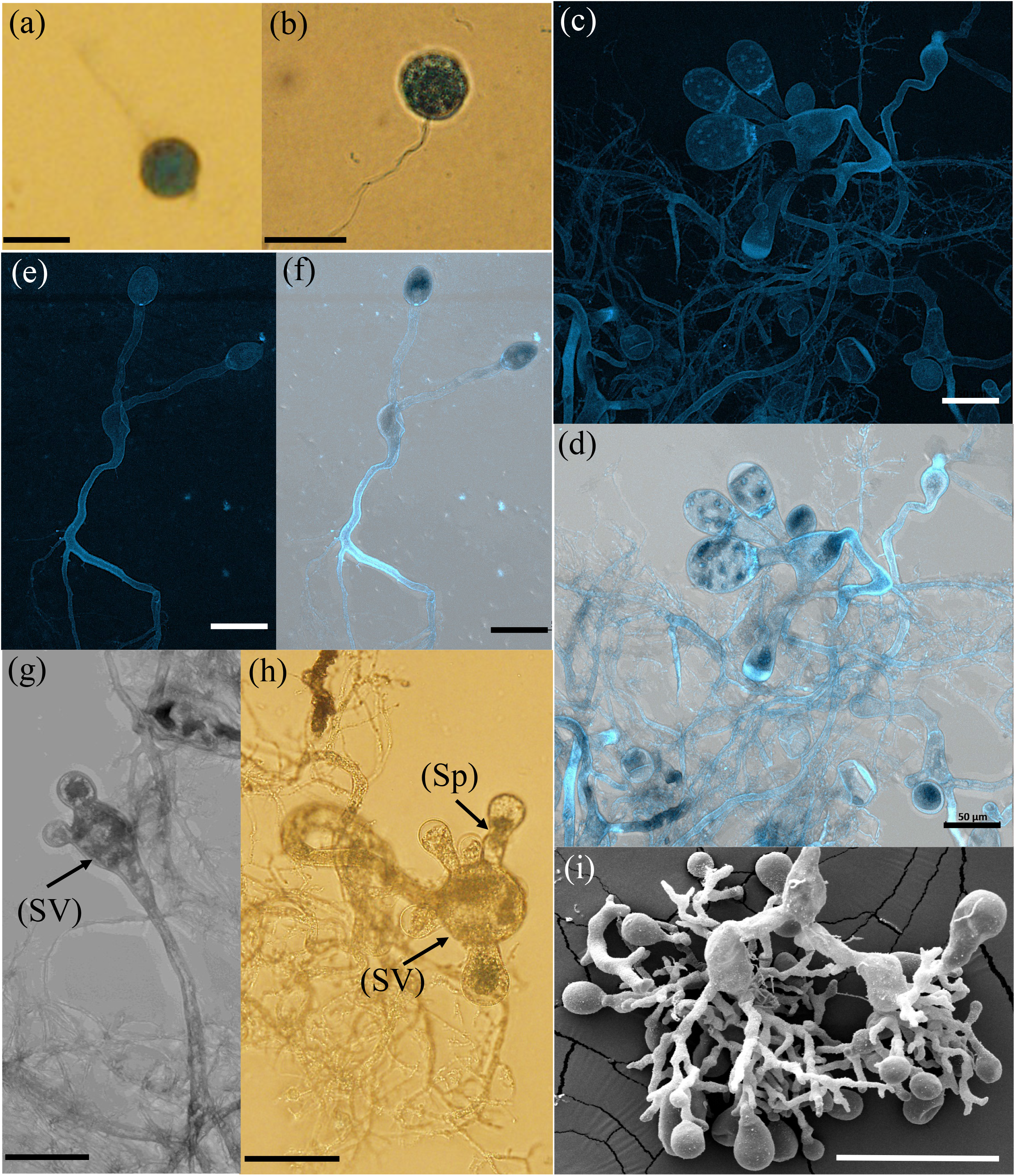
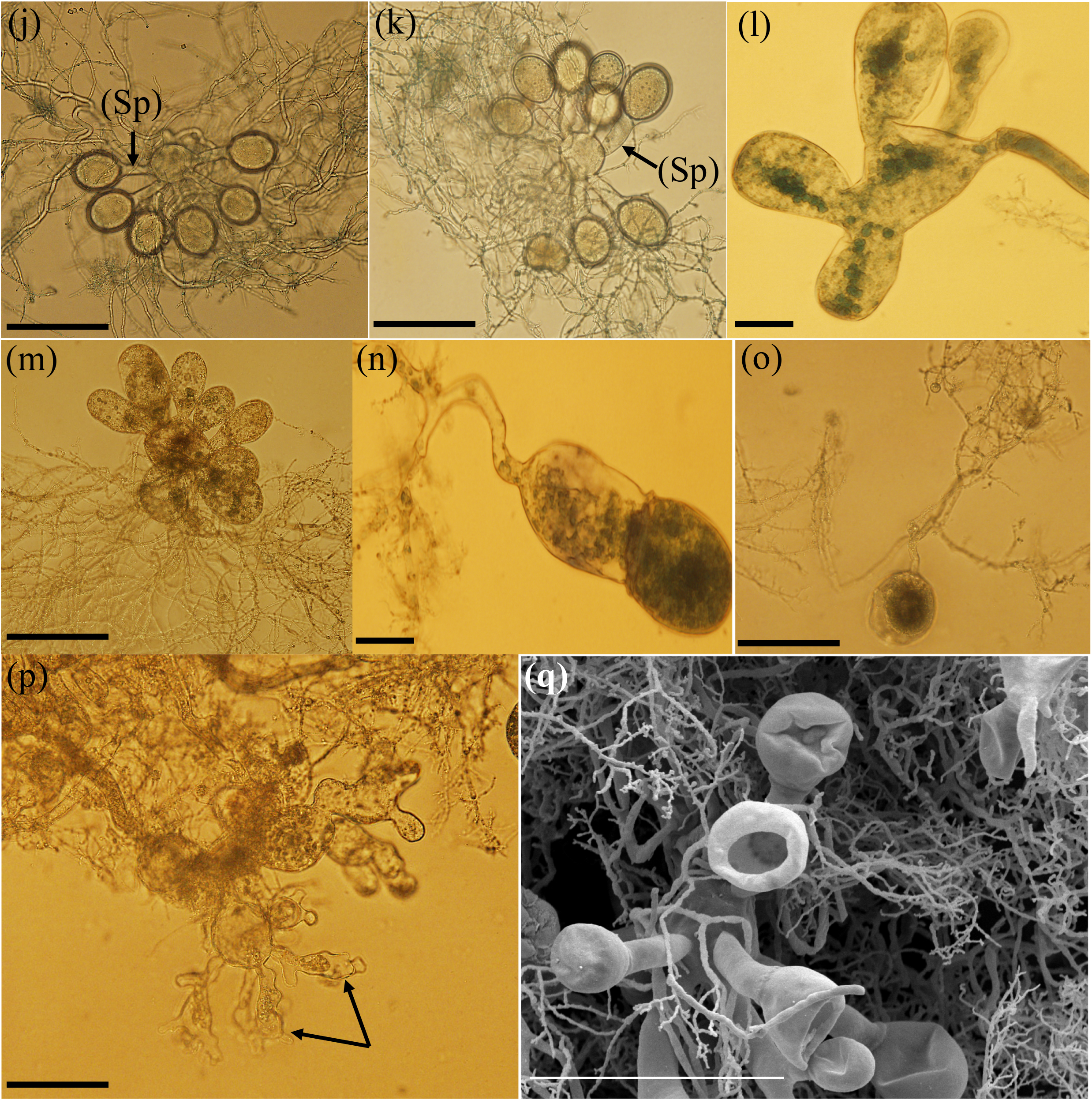
Microscopic features of *Paucimyces polynucleatus* strain BB-3. Light (a, b, h, and j-p), confocal (c-g), and scanning electron (i and q) micrographs are shown. Overlay images are shown in d and f. (a) A monoflagellated zoospore. (b) A biflagellated zoospore. (c-f) Polycentric thalli, with nuclei present in the rhizomycelim. Note the nucleated zoospores inside the mature sporangia (arrows). (g) Early thallus development stage starts by swelling of the hyphal tip forming a spherical vesicle (SV) and developing immature sporangia (S) (arrows). (h) Multiple sporangiophores (Sp) develop on the spherical vesicle (SV). (i) A mature thallus with multiple sporangia. (j-k) Ovoid sporangia developing at apices of sporangiophores (Sp), note the basal wall separating mature sporangia from the sporangiophores (arrows). (l-m) Sporangia developing directly on the spherical vesicles (SV). (n-o) Mature thalli with single ovoid sporangium. (p) An old culture producing empty sporangiophore initials (arrows). (q) An empty sporangium after zoospores release through a wide apical pore, with sporangial wall staying intact. Bar: a, b, and l= 20 μm, c-h, j,k, m-p= 50 μm, i and q= 100 μm

### Substrate utilization

Strain BB-3 was able to utilize a wide range of substrates as the sole carbon and energy source (Table 1). The monosaccharides glucose, xylose, mannose, and fructose all supported growth, whereas glucuronic acid, arabinose, ribose, and galactose failed to sustain the viability of strain BB-3 cultures. Strain BB-3 was able to utilize all disaccharides tested including cellobiose, sucrose, maltose, trehalose, and lactose. Out of the polymers tested, strain BB-3 was able to grow only on cellulose, xylan, starch, and raffinose, but not inulin, polygalacturonate, chitin, alginate, pectin, peptone, and tryptone (Table 1).

### Phylogenetic analysis and ecological distribution

The D1/D2 regions of strain BB-3 showed very low intra-strain sequence divergence (0-0.25%), and length (778-780 bp) heterogenicity. Similarly, the ITS1 region of strain BB-3 showed low intra strain sequence divergences (0-0.38%), and length (263-264 bp) heterogenicity. The ITS1 and D1/D2-LSU regions from strain BB-3 were 100% similar to sequences assigned to the uncultured lineage RH5 obtained in a previous culture-independent diversity survey from the same sample on which isolation was conducted (blackbuck deer), as well as few other samples (aoudad sheep, domesticated sheep, and Axis deer), demonstrating that these newly obtained isolates are cultured representatives of the RH5 lineage (Hanafy et al., 2020a).

In D1/D2 LSU trees, strain BB-3 formed a distinct cluster, within a broader supra-genus clade comprising the genera *Orpinomyces, Pecoramyces, Ghazallomyces, Neocallimastix, Feramyces*, and *Aestipascuomyces* (Figure 3a). D1/D2 LSU sequence divergences between strain BB-3 and its closest relatives in these lineages were 93.5% to *Orpinomyces joyonii* strain D3A, 91.05% to *Pecoramyces ruminantium* strain S4B, 92.32% to *Ghazallomyces constrictus* strain AXS31, 92.6% to *Neocallimastix cameroonii* strain G3, 91.2% to *Feramyces austinii* strain DS10, and 89.43% to *Aestipascuomyces dupliciliberans* strain A252. In ITS1 trees, the closest relatives were members of the genera *Orpinomyces, Pecoramyces, Ghazallomyces, Neocallimastix, Feramyces, Aestipascuomyces, Joblinomyces*, and *Agriosomyces* (Figure 3b). The closest cultured representative based on ITS1 sequence similarity was *Joblinomyces apicalis* strains GFH681 and SFH683 (81.25% similarity). Interestingly, strain BB-3 ITS1 sequence showed 95.2% similarity to an isolate described as *Anaeromyces* sp. strain W-98 (GenBank accession number AY091485), but no publication or documentation on the fate of that isolate is available.

**Figure 3.**
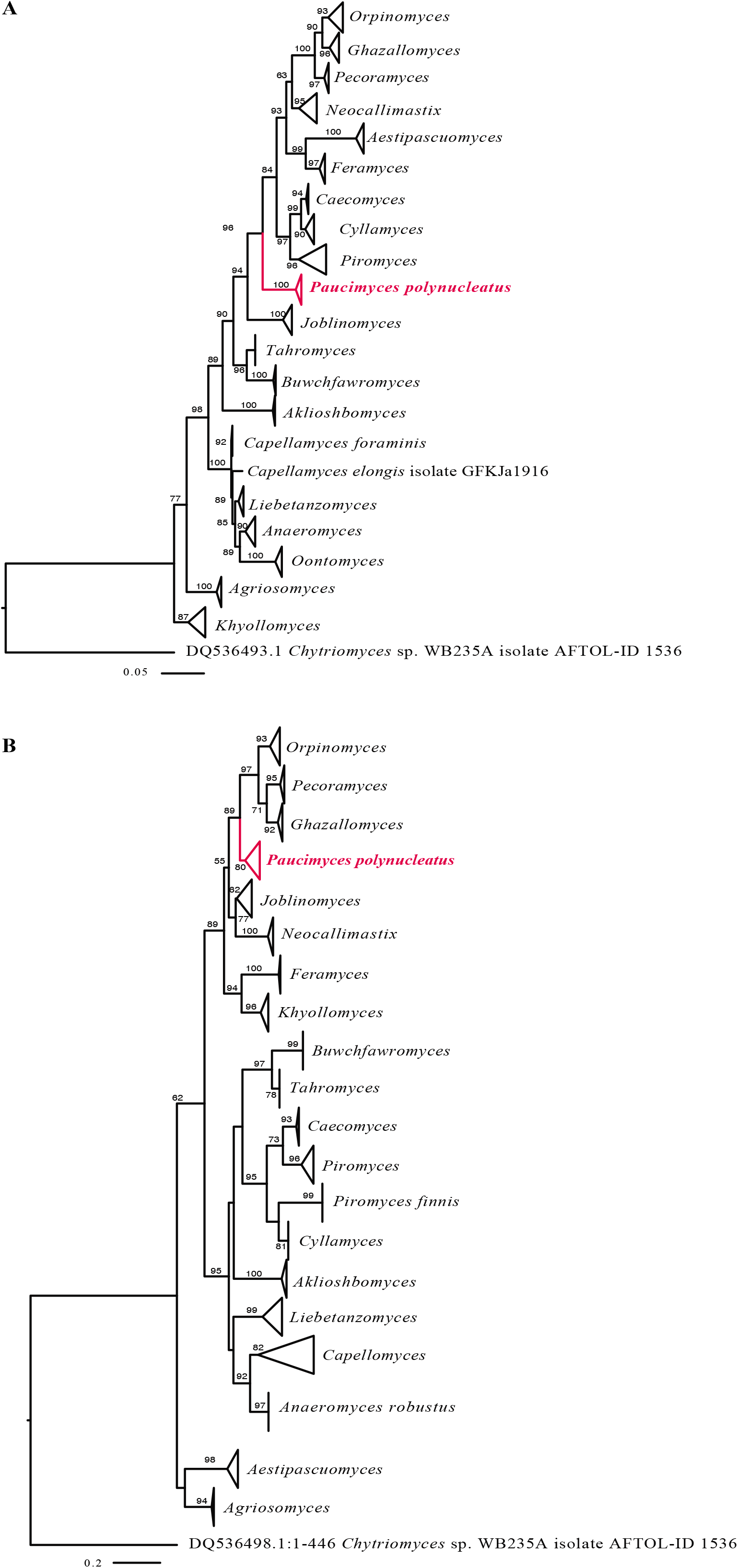
Phylogenetic affiliation of the *Paucimyces* clade to other AGF genera based on the sequences of (A) D1–D2 LSU and (B) ITS-1 sequences. Sequences were aligned in MAFFT (Katoh et al., 2019) and the alignments were used to construct maximum likelihood trees in FastTree using the GTR model. using *Chytriomyces* sp. WB235A isolate AFTOL-ID 1536 was used as the outgroup. Bootstrap values (from 100 replicates) are shown for nodes with more than 50% bootstrap support.

## Discussion

Strain BB-3 represents the first cultured representative of the RH5 lineage, and would constitute the twentieth described genus within the phylum Neocallimastigomycota. In addition to its distinct phylogenetic position in AGF D1/D2 LSU and ITS1 trees (Figure 3a & b), strain BB-3 possesses multiple unique morphological and microscopic characteristics that differentiates it from all described AGF genera. Strain BB-3 exhibits a polycentric thallus growth pattern, in which the zoospore contents completely migrate into the germ tube that eventually develops into a nucleated rhizomycelium capable of producing multiple sporangia per thallus. This thallus development pattern has been encountered only in the AGF genera *Anaeromyces* (Breton et al., 1990), *Orpinomyces* (Barr et al., 1989), and *Cyllamyces* (Ozkose et al., 2001). However, there are several key morphological and microscopic features that clearly differentiate strain BB-3 from other polycentric AGF genera. For example, strain BB-3 has a filamentous rhizomycelium, distinguishing it from the characteristic bulbous rhizomycelium of the genus *Cyllamyces*. Compared to *Orpinomyces* spp., strain BB-3 exhibits a thin and loose biofilm-like growth in liquid media, and produces small compact colonies (2-4 mm), unlike the cottony growth pattern and the large (usually >1 cm diam.) colonies characteristic of *Orpinomyces* spp. Microscopically, strain BB-3 produces monoflagellated, occasionally biflagellated, zoospores in contrast to the polyflagellated *Orpinomyces* zoospores. Compared to *Anaeromyces* spp., strain BB-3 produces a non-constricted hyphae and ovoid sporangia, unlike members of *Anaeromyces* spp. that are known to produce constricted sausage-shaped hyphae and mucronate sporangia.

A diagnostic characteristic of strain BB-3 is the formation of spherical vesicles (swellings at the hyphal tips) (Figure 2g-h) from which multiple sporangia are formed either directly on the spherical vesicles (Figure 2l-m) or at the end of a sporangiophore (Figure 2j-k). Such feature has rarely been observed in previously-reported taxa. Notably, a single isolate designated *Piromyces polycephalus* and isolated from the rumen fluid of water buffalo, was found to display a similar sporangial development pattern (Figures 3 and 4 in (Chen YC, 2002)). The proposed affiliation with the genus *Piromyces* implies a monocentric growth pattern, although the pictures do not clearly show the growth pattern and nuclear localization. Unfortunately, the absence of extant culture of *P. polycephalus* prevents further investigation into this issue. Also, lack of sequence data for *P. polycephalus* precluded our full understanding of the phylogenetic relationship between *P. polycephalus* and strain BB-3.

D1/D2 LSU sequences representing lineage RH5 were identified in one recent culture-independent diversity survey, where it was encountered in 10/31 fecal animal samples, and constituted >10% in only two samples (a domesticated sheep and a wild blackbuck antelope) (Hanafy et al., 2020a). RH5 sequences were identified in foregut fermenters (9 out of 10 samples), with a 0.2% relative abundance in miniature Donkey samples, the sole hindgut animal that harbored this lineage. ITS1 sequences similar to the RH5 lineage were also identified in four zoo-housed animals including American Bison, Llama, Sable Antelope, and Western tufted deer with a relative abundance of 0.03%, 1.2%, 18.93%, and 0.03% respectively. These sequences originated from a previous culture-independent survey conducted on zoo-housed animals (Liggenstoffer et al., 2010). Collectively, this pattern suggests a limited global distribution of lineage RH5 in the herbivorous gut, and a clear preference to ruminants over hindgut fermenters. However, studies on anaerobic gut fungal diversity are relatively sparse, localized, and lack spatiotemporal dimensions. As such, these observations should be regarded as preliminary, and more in-depth sampling and diversity assessment efforts are needed to confirm, disprove, or identify additional patterns governing the distribution of this lineage.

Thallus development pattern is a key feature used in the classification of the basal fungal lineages including the Neocallimastigomycota. Strain BB-3 exhibited a classical polycentric thallus growth, i.e. multiple sporangia per thallus. This growth pattern is associated with migration of the nucleus out of the zoospore into the germ tube, which elongates and branches into rhizomycelium. Within the rhizomycelium, repeated nuclear divisions occur and nuclei migrate into individual hyphae, resulting in a fungal thallus of unlimited extent and with multiple sporangia. Such pattern is in contrast to the monocentric thallus growth (single sporangium per thallus), where the rhizoid is devoid of nuclei and the thallus is of determinate extent with a single sporangium (Hess et al., 2020; Ho and Barr, 1995). It is worth noting that the presence of multiple sporangia per thallus is a hallmark of polycentric growth. However, some monocentric genera such as *Caecomyces communis, Piromyces polycephalus, Khyollomyces ramosus* produce branched sporangiophores with two or more sporangia resulting in a multi-sporangiate thallus (Chen YC, 2002; Hanafy et al., 2020b; Ho and Barr, 1995). In addition to the Neocallimastigomycota (Ho and Barr, 1995), polycentric growth pattern is known to occur in several basal fungal lineages, e.g. the genera *Nowakowskiella* and *Cladochytrium* in the phylum Chytridiomycota (Barr, 1978). Phylogenetic analysis shows that polycentric genera are polyphyletic within the Neocallimastigomycota, suggesting that multiple events of acquisition/loss of this trait has occurred throughout Neocallimastigomycota evolution and obscuring the nature of the AGF last common ancestor. The genetic and epigenetic determinants of this phenotypic pattern is yet unclear, hindered by the absence of genome representatives from most of the currently described AGF genera (Solomon et al., 2016; Youssef et al., 2013). Similarly, the niche preference of polycentric versus monocentric taxa, and correlation between such growth pattern and ecological distribution is murky. It is notable that prior studies have suggested that all previously described polycentric genera (*Anaeromyces, Orpinomyces*, and *Cyllamyces*) appear to exhibit a distribution pattern where they are present in the majority of examined animals, but often in low relative abundance. In contrast, RH5 appear to have a much more limited distribution, but could represent a majority of the community in rare cases (Hanafy et al., 2020a; Liggenstoffer et al., 2010).

Based on morphological, physiological, microscopic, and phylogenetic characteristics, we propose accommodating these new isolates into a new genus, for which the name *Paucimyces polynucleatus* is proposed. The type strain is *Paucimyces polynucleatus* strain BB-3.

## TAXONOMY

### Paucimyces

Radwa A. Hanafy, Noha H. Youssef, and Mostafa Elshahed, gen. nov. Mycobank accession number: MB838953

#### Typification

*Paucimyces polynucleatus* Radwa A. Hanafy, Noha H. Youssef, and Mostafa Elshahed

#### Etymology

*Pauci* = derived from the Latin word for few, reflecting its relatively limited distribution in nature; *myces* = the Greek name for fungus.

Obligate anaerobic fungus that produces polycentric thallus with highly branched nucleated rhizomycelium of indeterminate length. The fungus is characterized by formation of spherical vesicles at the hyphal tips. Multiple sporangia are developed either directly on the spherical vesicles or the end of sporangiophores. Mature sporangia are separated from the sporangiophores by basal walls. Old cultures produce sporangiophore initials with no sporangia. Zoospores are mainly monoflagellated. Bi-flagellated zoospores are occasionally encountered. Frequent sub-culturing results in cultures that lose the zoosporogenesis ability and produce sterile sporangia. The clade is defined by the sequence MW694896 (for ITS1, 5.8S rDNA, ITS2, D1-D2 28S rDNA). The most genetically similar genera are *Orpinomyces*, which is characterized by its polyflagellated zoospores and polycentric thallus that produce sporangia that are either terminal or intercalary, and *Joilinomyces*, which is defined as producing monocentric thalli and monoflagellated zoospores.

### Paucimyces polynucleatus

Radwa A. Hanafy, Noha H. Youssef, and Mostafa Elshahed Mycobank accession number: MB838954.

#### Typification

The holotype (Figure 2c) was derived from the following: U.S.A. OKLAHOMA: Stillwater, 36.12°N, 97.06°W, ∼300 m above sea level, 3-d old culture, isolated from frozen fecal samples of a wild blackbuck antelope (*Antilope cervicapra*) in December 2020 by Radwa Hanafy. Ex-type culture BB-3 is stored on solid agar media at 39°C at Oklahoma State University. GenBank accession number MW694896 (for ITS1, 5.8S rDNA, ITS2, D1-D2 28S rDNA).

#### Etymology

The species epithet (***polynucleatus***) reflects the polynucleated filamentous rhizomycelium produced during growth.

An obligate anaerobic fungus that produces globose (6-10 μm in diameter) monoflagellated zoospores. Biflagellated zoospores are occasionally observed. Flagellum length ranges from 15-30 μm. Zoospores germinate into polycentric thalli with extensively branched nucleated rhizomycelium of indeterminate extent. Spherical vesicles are developed at the hyphal tips, and multiple sporangia are developed directly on the spherical vesicles or at the end of sporangiophores. Sporangia are mainly ovoid and ranged in size between (15–90 μm L) X (10– 55 μm W). Old cultures produce empty sporangiophores initials. Also, prolonged sub-culturing results in sterile sporangia that fail to differentiate into zoospores. Cultures grown in cellobiose liquid media exhibit a thin loose biofilm-like growth and form white compact circular filamentous colonies (2-4 mm diameter) on agar roll tubes. The clade is defined by the sequence MW694896 (for ITS1, 5.8S rDNA, ITS2, D1-D2 28S rDNA).

#### Additional specimens examined

U.S.A. OKLAHOMA: Stillwater, 36.12°N, 97.06°W at ∼300 m above sea level, isolated from frozen fecal samples of a wild blackbuck antelope (*Antilope cervicapra*), in December 2020 by Radwa Hanafy. These cultures are named BB-2, BB-12, and BB-14.

## Acknowledgments

This work has been supported by NSF grant 2029478 to NHY and MSE.

## Notes

### Competing Interest Statement

The authors have declared no competing interest.

